# Optimized Ex Vivo Differentiation of CD103^+^ Dendritic Cells and High-Efficiency Retroviral Transduction of Mouse Bone Marrow HSCs

**DOI:** 10.1101/2025.09.21.677651

**Authors:** Mukta Asnani, Jiekun Yang

**Affiliations:** Department of Genetics, School of Arts and Sciences, Rutgers University-New Brunswick, Piscataway, NJ, USA; Human Genetics Institute of New Jersey, Rutgers University-New Brunswick, Piscataway, NJ, USA

## Abstract

CD103^+^ conventional dendritic cells (cDC1s) are key drivers of antitumor immunity, but their scarcity and resistance to genetic manipulation make them difficult to study. We optimized a two-stage ex vivo culture system using key cytokines and growth factors to efficiently generate CD103^+^ cDC1-like cells from mouse bone marrow progenitors. These cells closely mimicked their in-vivo counterparts, displaying CD103 expression, robust cytokine production, and functional responses to immune stimulation. Additionally, we established a high-efficiency retroviral transduction method using ecotropic pseudotyped virus and retronectin-coated plates, significantly improving gene delivery into mouse hematopoietic stem cells. This integrated platform provides a powerful approach for dissecting CD103^+^ cDC1 biology and advancing dendritic cell–based immunotherapy research.

## Introduction and Results

Dendritic cells (DCs) play a central role in regulating immune responses, balancing activation and tolerance through their ability to cross-present antigens and coordinate antigen-specific adaptive immunity [1, 2]. As potent antigen-presenting cells (APCs) of the innate immune system, DCs bridge innate and adaptive immunity in cancer and infectious diseases. Their development begins in the bone marrow, where progenitors committed to the conventional DC (cDC) lineage (pre-DCs) exit the bone marrow and migrate to peripheral tissues [3, 4]. There, they complete terminal differentiation and maturation upon encountering and phagocytosing antigens, including tumor-associated antigens. Mature DCs then cross-present these antigens via MHC class I or II complexes and migrate to draining lymph nodes, where they activate naïve T cells, driving their proliferation, activation, and differentiation [5, 6].

Dendritic cells can be broadly classified into three major subsets: classical DCs (cDCs), plasmacytoid DCs (pDCs), and monocyte-derived DCs (moDCs), each defined by distinct ontogeny, surface marker expression, and functional specialization [7]. In mice, cDCs are further divided into two main subsets: CD103^+^ cDC1s, which primarily reside in non-lymphoid tissues, and CD8α^+^ cDC2s, which are enriched in lymphoid tissues [8]. While all DC subsets are capable of initiating T-cell immune responses, CD103^+^ cDC1s are particularly adept at surveying peripheral tissues, undergoing maturation, migrating to draining lymph nodes, and cross-presenting antigens to efficiently prime naïve CD8^+^ T cells [9-12]. Recent advances in next-generation sequencing technologies, including single-cell and spatial transcriptomics, have uncovered previously unrecognized heterogeneity within DC populations. These studies have identified additional subpopulations beyond cDC1 and cDC2, including a distinct cluster of mature DCs—termed mregDCs or DC3s—that have been described across multiple human cancers. mregDCs are characterized by elevated expression of costimulatory molecules (CD40, CD80, CD86), immunoregulatory markers (CCR7, LAMP3), and high levels of MHC class II molecules, reflecting their potent antigen-presenting capacity. Functionally, mregDCs produce key cytokines such as IL-12p40/p70 and are highly effective at polarizing and activating CD8^+^ T cells.

The activation and functional programming of these DC subsets are largely shaped by pattern recognition receptors, such as Toll-like receptors (TLRs), which detect pathogen-associated molecular patterns (PAMPs) [2]. The repertoire of TLRs expressed by each DC subset enables them to respond to diverse stimuli, engage distinct signaling pathways, and contribute to the functional heterogeneity observed within the tumor microenvironment.

The presence of mregDCs within the tumor microenvironment (TME) has emerged as a critical factor in shaping anti-tumor immunity, with their abundance positively correlating with improved prognosis and better responses to immunotherapy across multiple solid tumor types [13]. However, the immunosuppressive nature of the TME can drive DC dysfunction, limiting their ability to mount effective immune responses [14]. High infiltration of immunosuppressive cell populations, combined with tumor-associated stressors such as hypoxia and lactate accumulation, can impair DC maturation, antigen uptake, and expression of key costimulatory molecules [15, 16]. These dysfunctional DCs fail to efficiently activate T cells, contributing to immune evasion and tumor progression.

Consequently, strategies aimed at restoring or enhancing DC activation and function are gaining significant interest as potential immunotherapeutic approaches. Investigating the molecular and cellular regulators that control the expression of DC stimulatory and immunoregulatory markers is therefore essential for understanding their biology within the TME and for designing interventions that reinvigorate anti-tumor immunity.

A major challenge in studying primary DCs—particularly cDC1s—is their scarcity in blood and tissues, which limits their availability for detailed analysis, functional studies, and therapeutic exploitation [17]. To overcome this limitation, researchers frequently rely on in vitro culture systems to generate larger numbers of cDCs. These systems aim to recapitulate the transcriptional, phenotypic, and functional characteristics of their in vivo counterparts, enabling more robust mechanistic studies and the exploration of DC-based immunotherapies.

Similar to other immune cell types, DCs can be generated in vitro from bone marrow progenitors, with Fms-like tyrosine kinase 3 ligand (Flt3L) serving as a key driver of cDC development. This was first demonstrated by studies showing that subcutaneous administration of Flt3L in mice leads to a marked expansion of cDC populations [18]. Over time, multiple in vitro differentiation protocols have been developed to generate DCs that closely resemble their in vivo counterparts at the transcriptomic and functional levels. One widely used approach involves a two-stage culture system in which hematopoietic stem cells (HSCs) are first expanded in the presence of stem cell factor (SCF or KitL) and then driven to differentiate into DC subsets with Flt3L [19]. KitL is thought to recruit and expand Flt3^−^ progenitors, thereby enhancing cDC1 output. However, the definition of cDC1s in these early studies was based primarily on CD24 expression rather than CD103 or CD8α, which are more specific markers for tissue-resident cDC1s. Because CD24 is also expressed by other immune cells such as macrophages, these findings may have overestimated true cDC1 generation.

Moreover, bone marrow cultures treated with Flt3L alone produce cDC1-like cells that express the lineage-defining transcription factor IRF8 but lack canonical cDC1 surface markers such as CD103, CD8α, and DEC-205 [20, 21], limiting their utility in dissecting CD103^+^ cDC1-specific biology. Co-culturing BM HSCs with Notch-ligand DLL1–expressing OP9 stromal cells yields DCs enriched for cDC1-associated markers, but these cells more closely resemble lymphoid or splenic cDC1s, characterized by high CD8α and low CD103 expression. Since splenic cDC1s naturally lack CD103, the Notch-ligand approach does not fully capture peripheral tissue–resident CD103^+^ cDC1 biology. Interestingly, although granulocyte-macrophage colony-stimulating factor (GM-CSF) is classically associated with the generation of monocyte-derived DCs (moDCs), it has also been shown to promote the proliferation and persistence of CD103^+^ DCs when combined with Flt3L in bone marrow cultures [22]. These in vitro–derived CD103^+^ DCs (iCD103-DCs) closely mimic the phenotype and functional properties of tissue-resident cDC1s, including efficient migration to draining lymph nodes and robust induction of T-cell–mediated immune responses. However, the potential synergistic effect of combining GM-CSF with KitL in this context remains to be fully explored.

To evaluate culture conditions that enhance in vitro cDC1 generation, we established multiple two-stage bone marrow culture systems (Fig. 1A). Hematopoietic stem and progenitor cells were isolated from the femurs and tibias of healthy mice and expanded in StemSpan SFEM medium supplemented with 10% FBS and various combinations of Flt3L, SCF (KitL), and GM-CSF at defined time points along the differentiation timeline. Cultures were monitored by flow cytometry at multiple stages to track the kinetics of key cDC subset markers, including CD24 and CD103 for cDC1s, and CD11b and CD172a for cDC2s.

**Figure 1.**
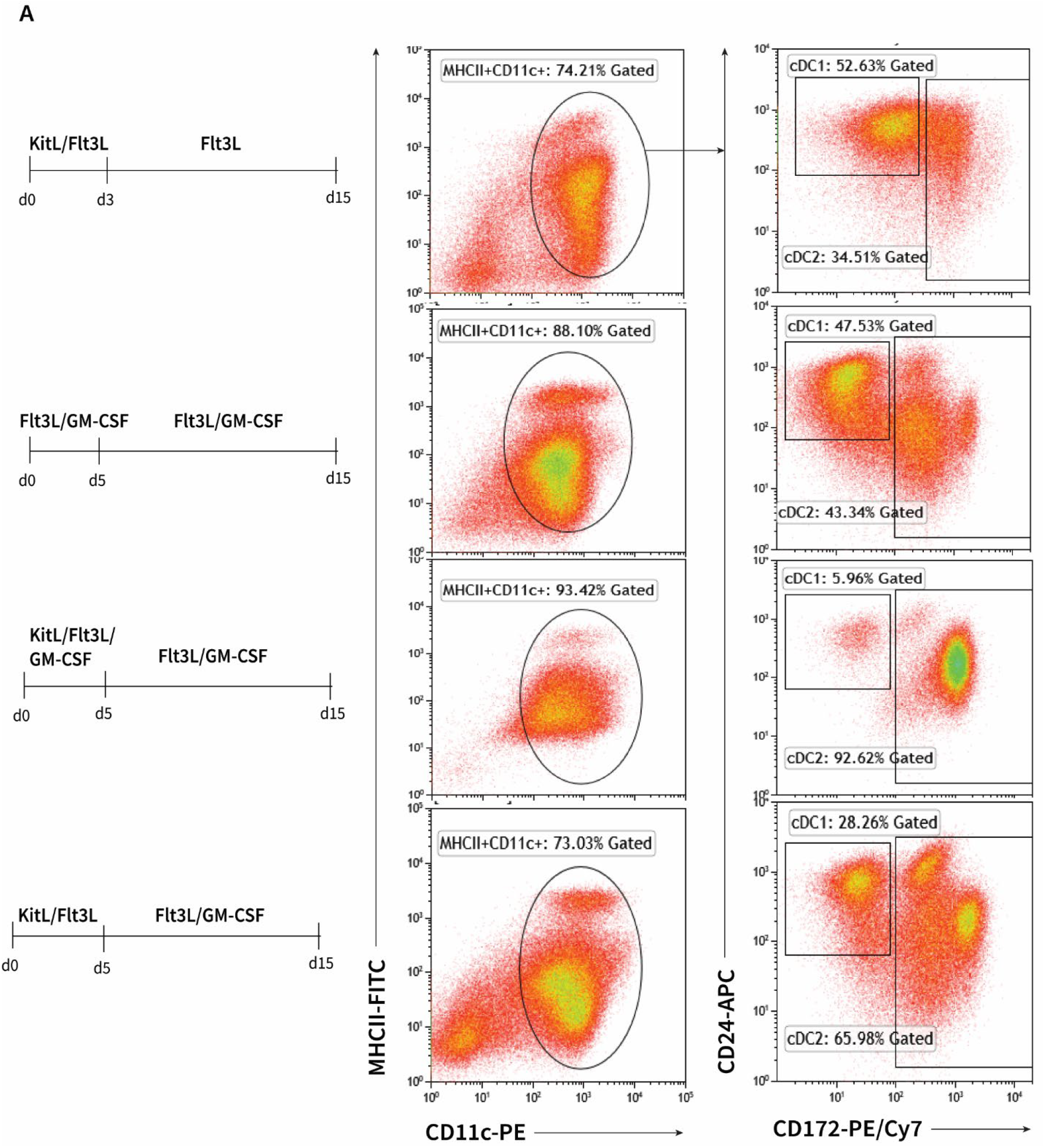
Ex vivo generation of cDCs from mouse bone marrow–derived HSCs. (A) Flow cytometry plots showing CD11c+MHCII+cells and the relative distribution of cDC subsets across different culture conditions.

Consistent with previous reports, the KitL/Flt3L culture system successfully generated CD24^+^/MHCII^+^/CD11c^+^ cDC1-like cells (Fig. 1A, top panel). In comparison, cultures supplemented with Flt3L and GM-CSF yielded a higher proportion of MHCII^+^/CD11c^+^ cells overall, with a corresponding increase in CD24^+^ cells (Fig. 1A, second panel). Interestingly, when KitL was added in combination with Flt3L and GM-CSF during the expansion phase, we observed a marked increase in MHCII^+^/CD11c^+^ cells but almost complete loss of CD24^+^ cDC1-like cells, resulting in cultures dominated by CD172a^+^ cDC2s (Fig. 1A, third panel).

We next modified the culture conditions by removing GM-CSF during the expansion phase and adding it only during the differentiation phase. This adjustment improved cell viability during the expansion phase and produced similar overall frequencies of MHCII^+^/CD11c^+^ cells as the Flt3L/GM-CSF system, though the CD24/CD172a distribution differed (Fig. 1A, bottom panel).

To further characterize these ex vivo–derived DCs, we assessed CD103 expression in KitL/Flt3L–Flt3L/GM-CSF and Flt3L/GM-CSF–derived cultures (Fig. 2A). Both systems generated CD103^+^ cells; however, their frequency was higher and peaked earlier (day 9) in the KitL/Flt3L–Flt3L/GM-CSF cultures compared to Flt3L/GM-CSF cultures. Notably, CD103^+^ cells declined rapidly and were nearly absent by day 15 in both systems, highlighting their relatively short lifespan in vitro. In our system, these DCs lacked CD8a expression (data not shown).

**Figure 2.**
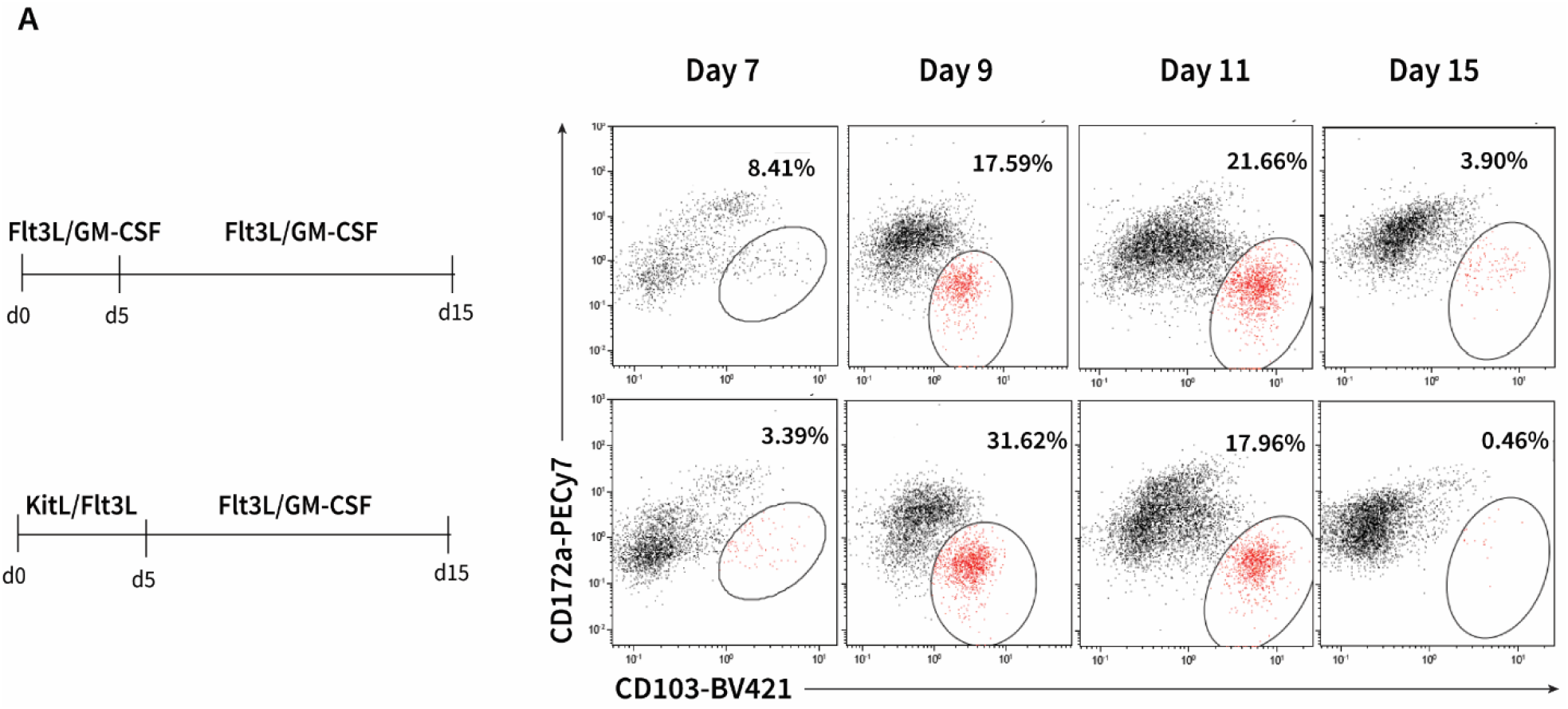
Time course analysis of CD103+ emergence in two different culture systems. (A) Flow cytometry plots showing emergence of CD103+ cDC1-like dendritic cells at different time points.

Next, we sought to determine whether ex vivo–derived cDCs from KitL/Flt3L–Flt3L/GM-CSF culture system, recapitulate the functional properties of in vivo cDC1s, particularly their capacity for cytokine production and PD-L1 (CD274) upregulation upon stimulation (Fig. 3A). Mouse cDC1s are known to express TLR3 as well as respond to cell-associated antigens via its expression of Clec9a [23, 24]. At day 9 of culture, stimulation with heat-killed *Listeria monocytogenes* (HKLM) led to robust upregulation of MHCII, PD-L1, and CD40 (Fig. 3B). Kinetic analyses revealed that these activation markers were significantly increased as early as 6 hours post-stimulation, coinciding with a subsequent near-complete loss of CD103^+^ cells by 24 hours.

**Figure 3.**
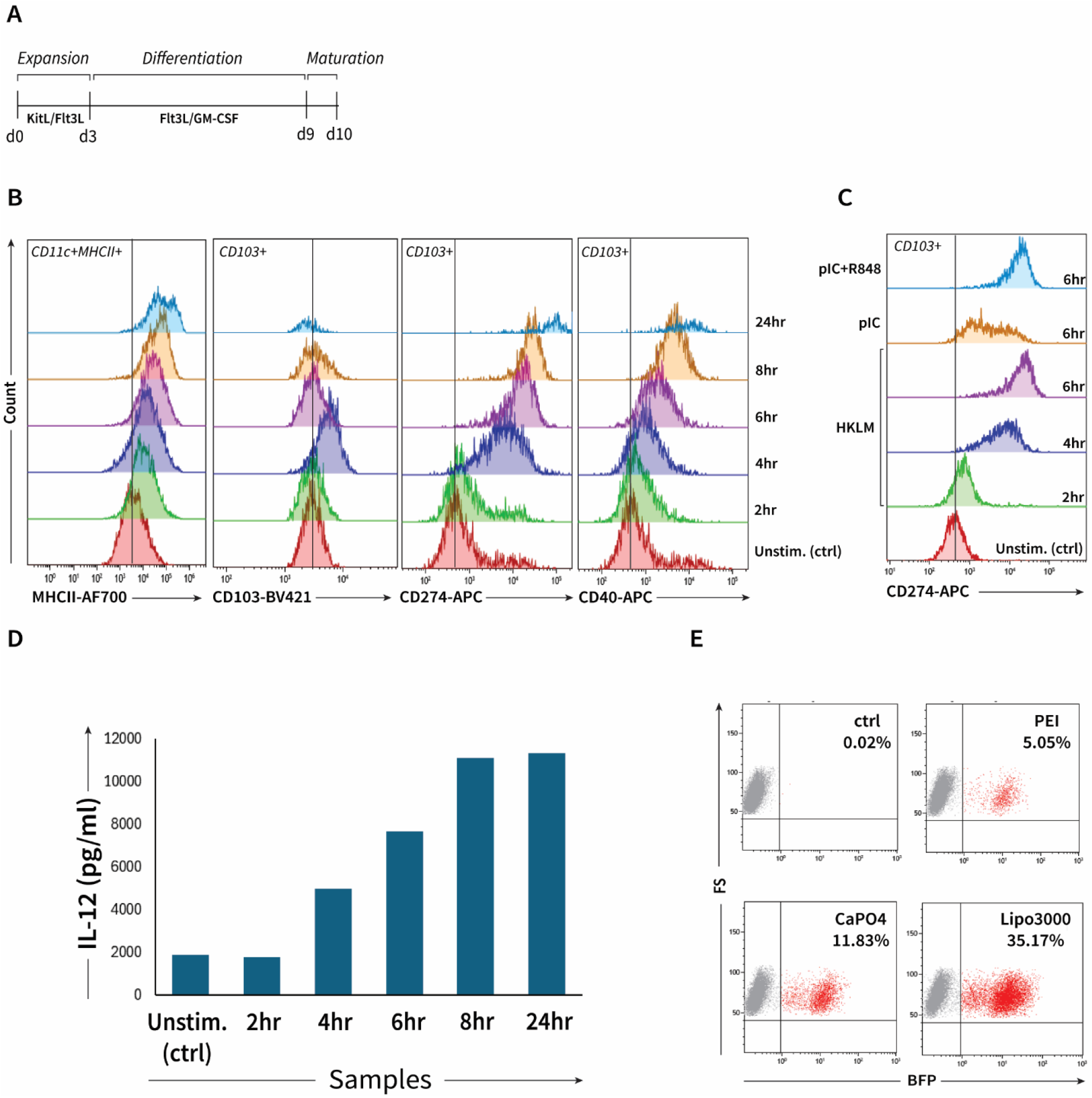
Functional characterization of ex-vivo derived CD103+ DCs. (A) Schematic of the culture method for BM expansion and differentiation. (B) Histograms of CD274 on CD103+ cells following incubation with TLR agonists and cell-associated antigen, HKLM. (C) Histograms of stimulation marker kinetics (MHCII, PD-L1/CD274, and CD40) on CD103+ cells following incubation with HKLM. (D) ELISA-based measurement of IL-12 secretion in supernatants from stimulated ex vivo cDC cultures. (E) Transfection efficiency of BM cells using Eco-pseudotyped retroviral BFP vectors generated with different transfection reagents.

Similarly, bone marrow–derived DCs responded to the TLR3 agonist poly(I:C), and this response was further enhanced by co-stimulation with the TLR7/8 agonist R848, consistent with previous studies (Fig. 3C). This activation correlated with strong IL-12 production, which was detected both in culture supernatants by ELISA (Fig. 3D). IL-12 levels peaked and plateauing around 8 hours, paralleling the kinetics of CD103^+^ cDC1 loss in culture. Interestingly, a transient increase in CD103^+^ cell numbers was observed at 4 hours, suggesting that IL-12 induction initially promotes cDC1 expansion but also accelerates terminal maturation and apoptosis, leading to the observed decline and complete loss of CD103^+^ cells by 24 hours (Fig. 3B).

Building on our characterization of ex vivo–derived DCs and their functional properties, we next sought to establish genetic tools to dissect the molecular pathways controlling their development and activation. A major hurdle in dissecting the biology of DCs is the limited ability to genetically manipulate these cells for functional studies. Both mouse hematopoietic progenitors and primary T cells are known to suppress lentiviral transgene expression in some contexts [25-27]. To overcome this, we tested alternative gene delivery approaches using a murine stem cell virus (MSCV)–based retroviral system, which has been shown to improve shRNA/sgRNA expression efficiency in primary murine cell types, particularly HSCs. Because VSV-G–pseudotyped lentiviral particles showed negligible infection rates in mouse bone marrow cells (data not shown), we utilized Plat-E packaging cells to generate retrovirus pseudotyped with ecotropic envelope proteins, which are highly permissive for murine cells.

To maximize viral titers, we compared multiple transfection reagents, including polyethylenimine (PEI), Lipofectamine 3000, and calcium phosphate, identifying the method that produced the highest infectious viral particles (Fig. 3E). Mouse HSCs were transduced on day 2 of culture during the KitL/Flt3L expansion phase. To further enhance transduction efficiency, we evaluated adjuvants that could promote infection without compromising cell viability. While polybrene was found to be toxic, retronectin-coated plates significantly increased transduction efficiency while maintaining viability.

Our findings suggest that ecotropic retroviral systems, combined with optimized transduction conditions such as retronectin coating, represent a robust platform for gene perturbation in murine HSC-derived DC cultures. This approach will enable functional genomic studies to identify key regulators of DC development, activation, and therapeutic potential.

## Conclusion

In this study, we systematically optimized the ex vivo generation, functional characterization, and genetic manipulation of murine cDC1s. We demonstrated that specific combinations of KitL, Flt3L, and GM-CSF drive robust expansion and differentiation of bone marrow–derived cDC1-like cells, with distinct effects on CD24, CD103, and CD172a expression. Functionally, these ex vivo–derived cDC1s recapitulate key features of their in vivo counterparts, including antigen responsiveness, PD-L1 upregulation, and IL-12 production, though they exhibit a relatively short lifespan. Finally, we established an efficient ecotropic retroviral transduction platform for genetic perturbation of murine HSC-derived DCs, paving the way for mechanistic studies and therapeutic applications. Together, these findings provide a scalable experimental framework for dissecting DC biology and for developing DC-based immunotherapies.

